# Being confident in confidence scores: calibration in deep learning models for camera trap image sequences

**DOI:** 10.1101/2023.11.10.566512

**Authors:** Gaspard Dussert, Simon Chamaillé-Jammes, Stéphane Dray, Vincent Miele

## Abstract

In ecological studies, machine learning models are increasingly being used for the automatic processing of camera trap images. Although this automation facilitates and accelerates the identification step, the results of these models may lack interpretability and their immediate applicability to ecological downstream tasks (e.g occupancy estimation) remain questionable. In particular, little is known about their calibration, a property that guarantees that confidence scores can be reliably interpreted as probabilities that a model’s predictions are true. Using a large and diverse European camera trap dataset, we investigate whether deep learning models for species classification in camera trap images are well calibrated, or in contrast over/under-confident. Additionally, as camera traps are often configured to take multiple photos of the same event, we also explore the calibration of predictions at the sequence level. Finally, we study the effect and the practicality of a post-hoc calibration method, i.e. temperature scaling, for predictions made at image and sequence levels. Based on five established models and three independent test sets, our findings show that, using the right methodology, it is possible to enhance the interpretability of the confidence scores, with clear implication for, for instance, the calculation of error rates or the selection of confidence score thresholds in ecological studies making use of artificial intelligence models.

## 1 Introduction

Camera traps have become a central tool in the monitoring and conservation of communities and populations. They generate a lot of data that can be used to infer, for instance, species richness, occupancy or activity patterns (Sollmann 2018). To exploit these data, it is first required to identify the species present in the photos or videos. This manual annotation task is generally long and tedious, but it has been shown in recent years that it can be replaced by an automatic classification made by artificial intelligence (AI; e.g. deep learning models), often with an accuracy of over 90% (Norouzzadeh et al. 2018; Willi et al. 2019; Whytock, Świeżewski, et al. 2021).

However, a responsible use of AI (Wearn et al. 2019) requires to understand whether results can be trusted or not, generally or per prediction. In a species classification model, accuracy (i.e., rate of true predictions) provides this information at the model’s level, whereas confidence scores should provide this at the prediction level. In many ecological studies, downstream tasks may directly rely on these scores, for instance when subsetting data considering that values above a certain threshold indicate true detections, or when propagating model uncertainty into subsequent statistical models. In these applications, confidence scores are frequently interpreted as probabilities of the predictions being true. However, it is often neglected (as probably unknown) that there is no guarantee that these scores can be interpreted in such way as many deep learning models may return biased confidence scores (Gawlikowski et al. 2023).

In the context of classification models, a model returning confidence scores that can be reliably interpreted as probabilities of the prediction being true is said to be well calibrated. For instance, if a model predicts the label of 100 images with a confidence score of 0.8, we would expect to observe an actual accuracy of 80% on these images. In the reverse, when this is not the case, a model is said to be under– or over-confident depending on the bias directionality. Whereas the question of calibration of confidence scores has been shown to be crucial in different fields such as autonomous driving (Bojarski et al. 2016) or medical diagnosis (Nair et al. 2018), it has rarely been studied or considered in ecological studies.

In the field of ecology, a poorly calibrated model (i.e. over– or under-confident) induces several issues. Indeed, as the score distribution is biased, confidence scores can not be interpreted as probabilities so that it is impossible to control for the accuracy (or error rate) associated to a given threshold value (Figure 1). In contrast, good calibration ensures the interpretability of the scores as probabilities allowing to control for the error rates in downstream tasks (Figure 1) such as occupancy estimation (Gimenez et al. 2022; Rhinehart et al. 2022), inference of species interaction (Parsons et al. 2022), realtime alert to guide law-enforcement (Whytock, Suijten, et al. 2023), confidence-based checking on citizen science platforms (e.g. Zooniverse (Simpson et al. 2014; Lotfian et al. 2021)) or confidence-based uploads to biodiversity inventories (August et al. 2020).

**Figure 1:**
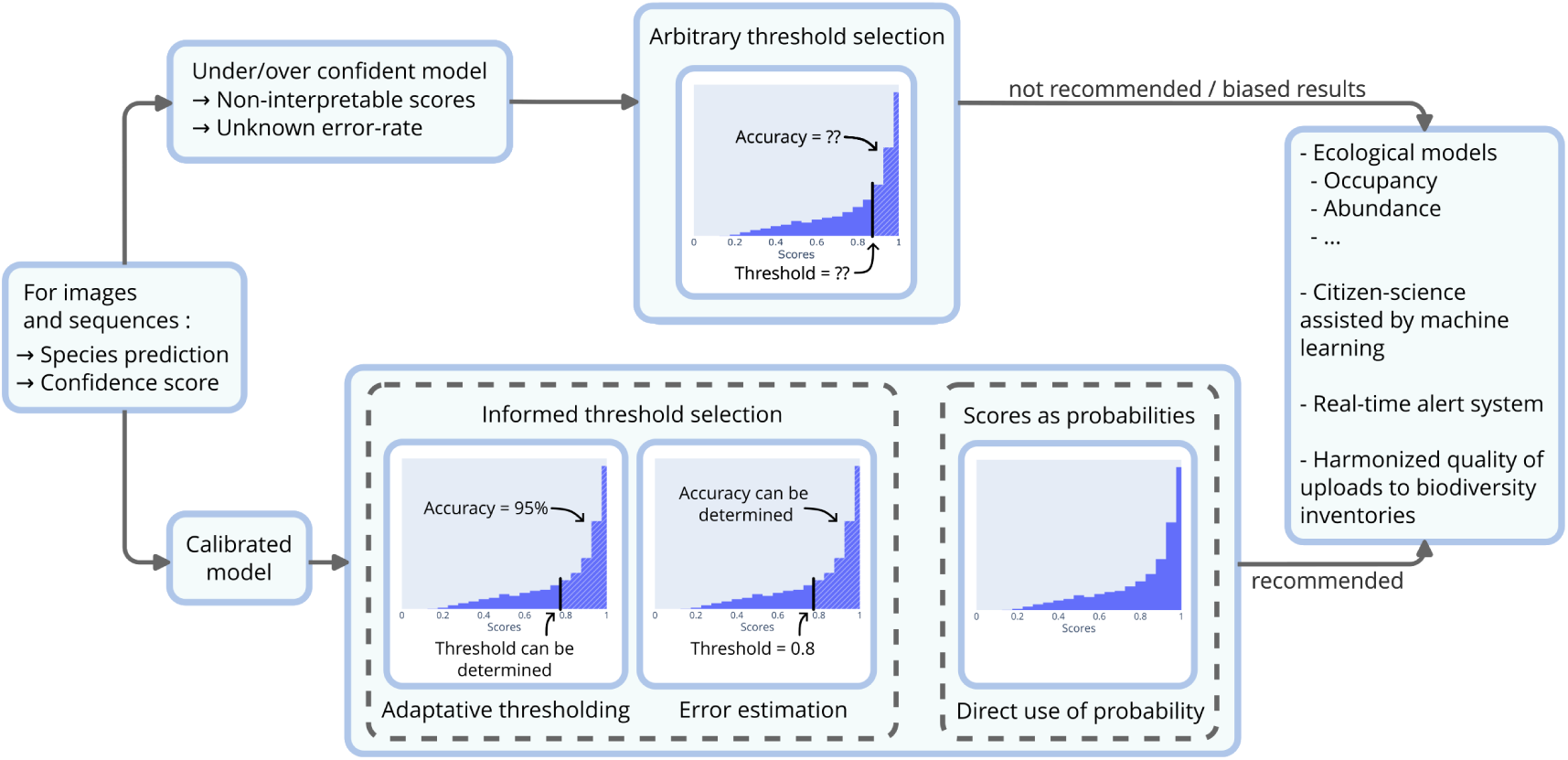
Practical use-cases where ecological downstream tasks (right) can be biased by the use of under/over-confident machine learning models (top), or enhanced by the use of calibrated models (bottom) when using deep learning species classification models for camera trap data (left).

In this paper, we explore the calibration of confidence scores in the context of species classification models for camera trap data. For that task, the recurring leading approach, as assessed in recent iWildcam competitions (Beery, Agarwal, et al. 2021), consists in two steps: (step 1) detecting animals, humans and vehicles and filtering out empty images using a robust detection model such as MegaDetector (Beery, Morris, et al. 2019; Mitterwallner et al. 2023) and (step 2) using a convolutional neural network (CNN) classification model to identify the species in the bounding box returned by the detection model, when an animal has been detected. We therefore focus on these species classification models (step 2), which are developed for a large range of species all over the world. We explore the interplay between accuracy and calibration for five state-of-the-art model architectures applied to camera trap data collected in three out-of-sample test data sets. Also, we consider the calibration of confidence scores at the level of sequences of images. Indeed, camera traps are often configured to take multiple photos at each trigger so that predictions are aggregated at the level of the sequence of images (sometimes called ‘the observation’ or ‘event’). The issue of the calibration of confidence scores at the level of sequences of images has not, to our knowledge, been addressed in the literature. Furthermore, we study the relevance of a popular post-hoc calibration method called temperature scaling (Platt 2000), for both image and sequence levels. Overall, on top of providing a solution to produce calibrated scores, our work intends to illustrate the benefits and effectiveness of calibration in downstream tasks with a practical use-case. Our findings show that we can estimate accurately the rate of classification errors made by a model and this important step can improve the pipeline of analyses based on camera trap data. Lastly, we use our results to provide a set of good practices for researchers and practitioners in the field.

## 2 Material and Methods

### 2.1 The DeepFaune Dataset

We use the dataset of the DeepFaune initiative (Rigoudy et al. 2023), which is a collaborative effort involving over 50 partners who, together, have gathered over two millions images and twenty thousand videos that they had manually annotated. These partners are affiliated to a wide range of institutions, such as organizations managing protected areas, hunting federations, and academic research groups. Images and videos were mainly collected in France, but also in a few European countries. Most of the annotation were at the species level, but some were at a higher taxonomic level (e.g. mustelid). Videos were converted into images by extracting frames of the first four seconds, with a time step of one second. The dataset provides a great diversity of habitats, elevations and weather conditions, as well as a wide variety of camera trap models with different settings, resolutions, flash type and image processing.

### 2.2 Training and validation datasets

For the species classification task, it is now well established that two-step approaches (object detector followed by a classifier) are more efficient than classifiers that process the whole image (Beery, Morris, et al. 2019; Norman et al. 2023). We use MegaDetector v5 (MdV5) (Beery, Morris, et al. 2019) to extract bounding boxes of animals, human and vehicles. Because MdV5 has already near-perfect accuracy on human and vehicles we only kept, for the training of our classifier, the bounding boxes that predicted the presence of an animal. For each bounding box, we created a cropped image of the original image, resulting in 429 347 cropped images of 22 different classes (the distribution of the classes is shown in Supporting Information Figure 1 and 2).

**Figure 2:**
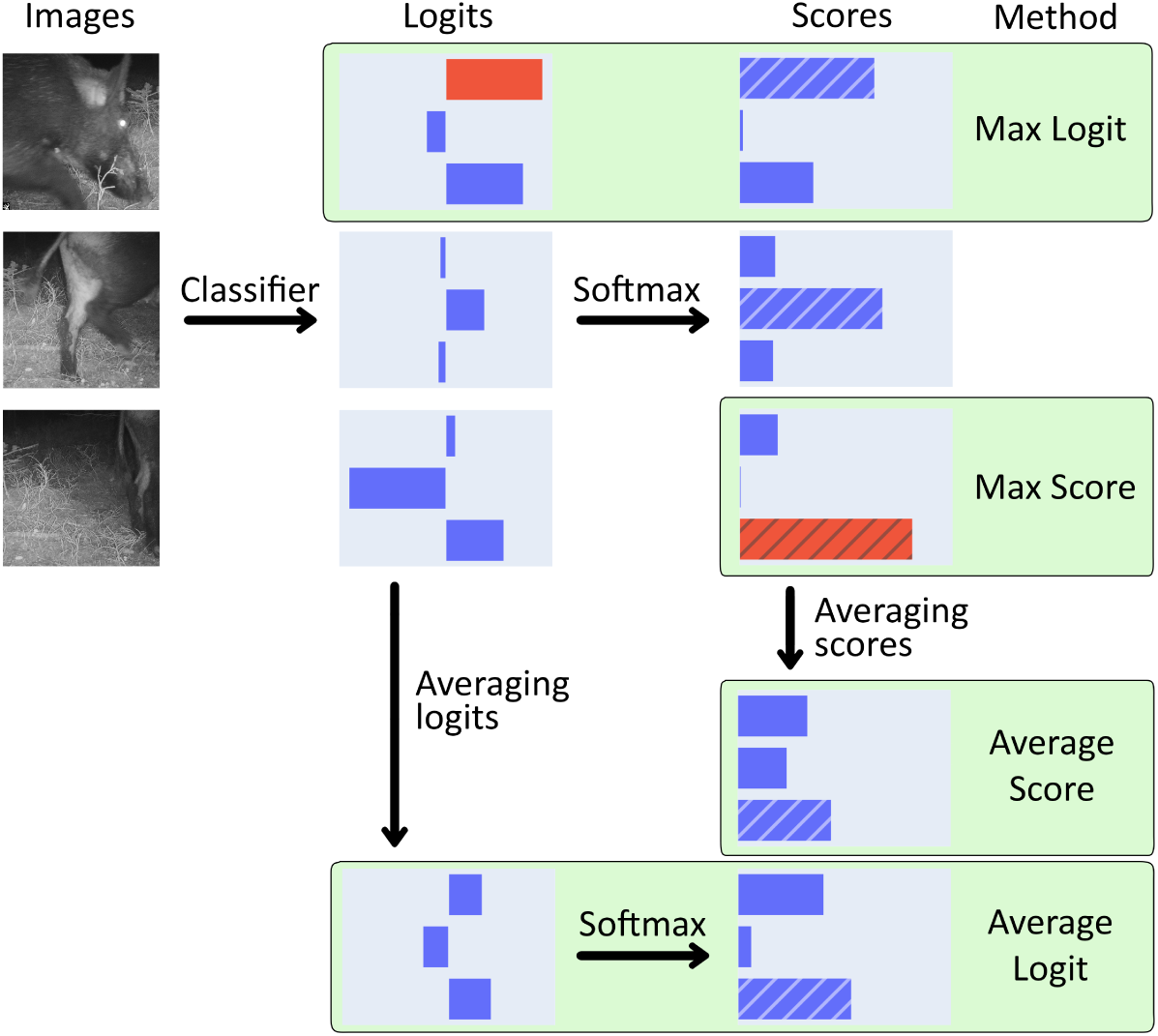
Illustration of the four aggregation methods. The greatest overall logit and score are in red. The top-1 score is hatched to emphasize that only this score is used to calculate the calibration.

To avoid overfitting and shortcut learning between the background of the images (i.e. camera trap site) and the observed species, we designed the training and validation sets to have disjoint pairs of background and species while having the same balance of species and diversity of habitats. The validation set represented about 20% of the images available while being disjoint from the training set at the species level: for each species, the validation set is made of images originating from partners different than the ones used in the training set, while being as close as possible to a 80/20 split. This requires solving a problem of combinatorial optimization known as *subset sum problem*, which is a special case of the *knapsack problem* and which can be achieved using dedicated libraries (e.g. mknapsack). Ultimately, we had 368 786 images in the training set and 60 561 in the validation set.

### 2.3 Out-of-sample test sets

To demonstrate that the results of the classifier could generalize beyond the images collected in the DeepFaune initiative, 3 out-of-sample test datasets were used. These datasets originated from ecological programs conducted in three geographically distinct areas. We refer to these datasets by the name of the areas they originate from:

- **Pyrenees**: camera trap study in the national reserve of Orlu in the French Pyrenees, conducted by the French Biodiversity Agency (OFB), 100 266 images and 12 species after preprocessing.
- **Alps**: camera trap study in the Ecrins national park in the French Alps, conducted by S. Chamailĺe-Jammes, 8 106 images and 12 species after preprocessing.
- **Portugal**: camera trap study in the Peneda-Ger^es National Park in Portugal (Zuleger et al. 2023), publicly available. 99 750 cropped images and 16 species after processing.

### 2.4 Sequences of images

It is common to configure camera traps to take a series of images after each trigger. It is therefore relevant to have a single prediction for the whole series of images. We thereafter name such series ‘sequences’. In our test sets, we considered that two consecutive images taken within 10s, at the same site (i.e. the same camera trap), belonged to the same sequence. We obtained sequences of 1 to 213 images.

### 2.5 Confidence score at sequence level

A sequence with *S* images has *S* individual predictions that can be aggregated in many different ways to produce a single prediction for the whole sequence. Formally, for a sequence of *S* images *x_i_*, the model predicts the logits 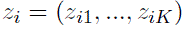 for each image, with *K* the number of classes. Confidence scores are derived using the softmax function: 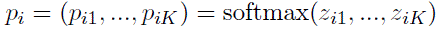. We aimed at predicting the confidence scores of the sequence 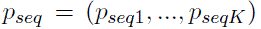 as a function of the predictions at the image level. We explored four different aggregation functions (Figure 2):

- **Average Score**: We averaged, over the sequence, the scores for individual pictures of the sequence: 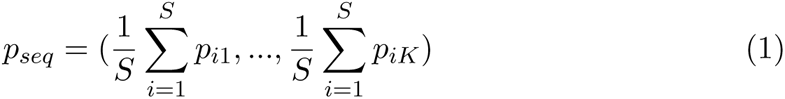
- **Average Logit**: We averaged, over the sequence, the logits for individual pictures of the sequence, and then applied the softmax function: 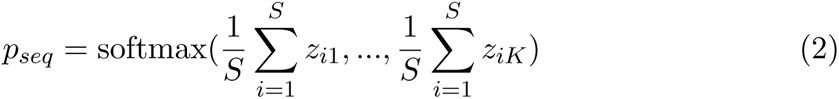
- **Max Score**: We kept the scores of the image that had the highest score amongst all scores of all images of the sequence: 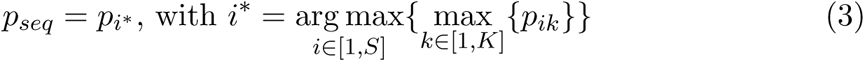
- **Max Logit**: We kept the scores of the image that had the highest logit amongst all logits of all images of the sequence: 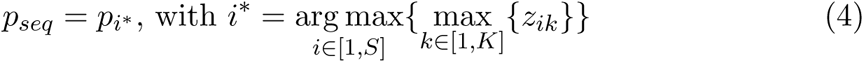

### 2.6 Calibration metrics

For a set of *N* images, we define the true class of the *i*-th image *y_i_* and *p_i_* = (*p_i_*_1_*, …, p_iK_*) the confidences scores of the *K* classes. The predicted class *y*^*_i_* is the top-1 classification prediction, that is the class with the greatest confidence score, denoted *s_i_*:

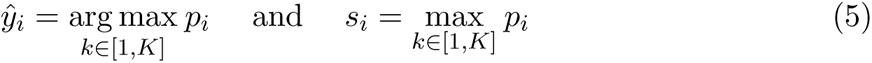

For *M* evenly spaced bins, we can define *b_m_* the set of indices *i* such as 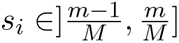 and compute the average bin accuracy and the average bin confidence score:

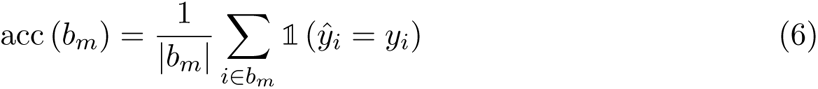

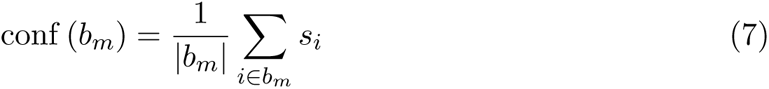

The bin-wise accuracy can be plotted to construct the reliability histogram (Guo et al. 2017) (e.g. Supporting Information Figure 3). It allows to visualize the calibration of a model: the closest the tops of the histogram bars are from the identity line, the better calibrated the model is. In addition, if the tops of the histogram bars are mostly above (resp. below) the line, the model is said to be under-confident (resp. over-confident).

**Figure 3:**
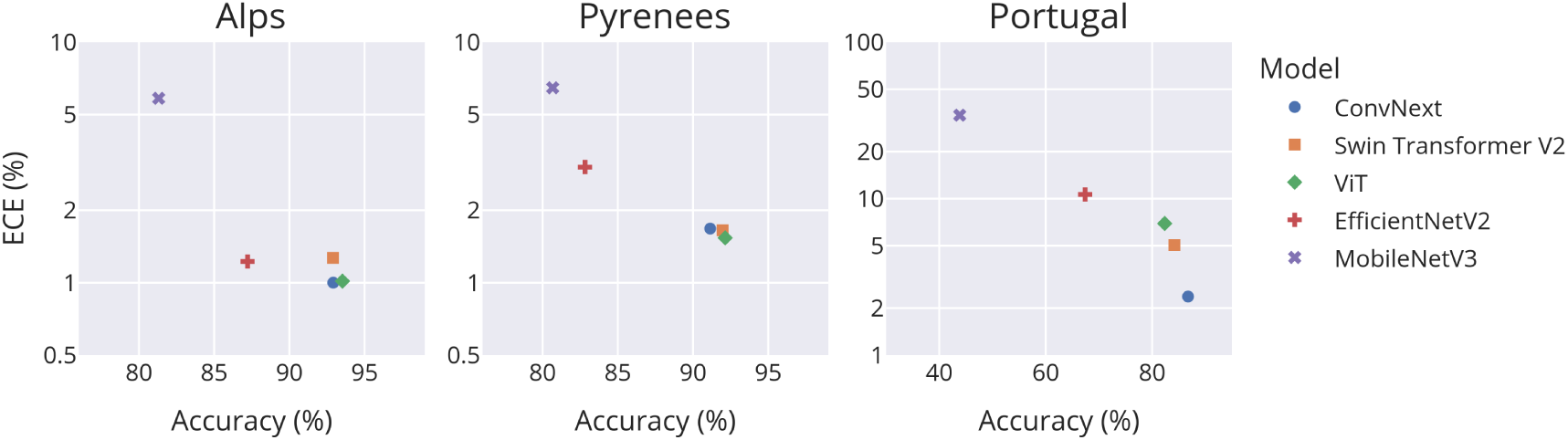
Scatterplot of ECE vs. accuracy values for five models (colored points) and three test data sets (panels), computed at the image level. Here, the ECE is not post-calibrated with temperature scaling (i.e. the temperature is 1 for all models).

The most common metric to measure the model’s calibration quantitatively is the Expected Calibration Error (ECE) (Guo et al. 2017). ECE is defined as the bin-wise calibration error weighted by the size of the bin:

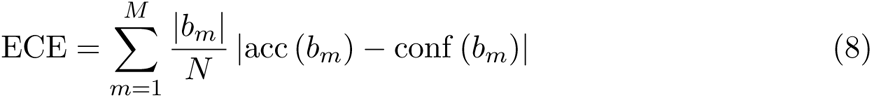

Due to the large amount of images in our test sets, we decided to use a greater number of bins, specifically 20 instead of the standard 15, to obtain a more precise measurement of calibration with the ECE. In addition to this metric, we evaluated the classification performance of our classifier with the accuracy metric. These two metrics can also be used to evaluate the classification and the calibration at the sequence level, using the score *p_seq_* and the associated predicted label 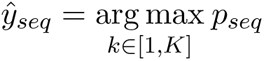.

### 2.7 Temperature Scaling

Temperature scaling (Platt 2000) is a post-processing method to improve the calibration of the model after the training. The scores predicted by the model are rescaled by a temperature parameter *T >* 0 using a generalization of the softmax function:

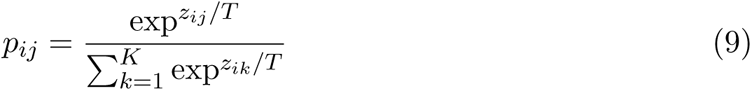

For *T* = 1 the scores obtained are the same as with the standard softmax function. *T >* 1 leads to lower scores and helps when the model is over-confident. Conversely, *T <* 1 increases the scores and helps under-confident models. For a given dataset, it is possible to determine the optimal temperature *T^∗^*, that minimize the ECE. However, this optimum temperature may differ from one dataset to another, and determining the optimum requires access to the labels. It is therefore unrealistic to use this individual temperature *T^∗^* to compare methods, as it cannot be calculated for a new dataset without manually annotating a fraction of the data. Instead, we propose to look at performance using a single temperature *T*^ˉ^ shared across the three datasets. We define *T*^ˉ^ as the temperature that minimizes the average ECE across the 3 test datasets. Temperature scaling can be combined with the four aggregation method (Section 2.5) to calibrate sequence level predictions by simply replacing the standard softmax function with Equation 9.

### 2.8 Deep learning models

To demonstrate the robustness of our findings, we used 5 established machine learning architectures: EfficientNetV2, ConvNext, ViT, Swin Transformer V2, and MobileNetV3. (Tan and Le 2021; Zhuang Liu et al. 2022; Dosovitskiy et al. 2021; Ze Liu et al. 2022; Howard et al. 2019). We have selected these architectures to represent CNNs (EfficientNetV2, ConvNext), Transformers (Swin, ViT), as well as lightweight architectures that could be deployed in camera traps that do the classification at the edge (MobileNetV3). Models were trained using the TIMM library (Wightman 2019) with transfer-learning from ImageNet-22k (Ridnik et al. 2021), the largest publicly available database. Data augmentation was applied using the imgaug library (A. B. Jung et al. 2020) using only standard transformations such as flips, crops, conversion to grayscale and affine transformation. The optimization was done using SGD, with a batch size of 32 and a different learning rate adapted for each architecture. To avoid overfitting, early stopping was used while monitoring the validation accuracy and with a patience of 10 epochs.

### 2.9 Error-rate estimation

For calibrated models, a predicted score is equivalent to the probability of the prediction being correct. Hence, for a given dataset analyzed with a classification model, the sum of the predicted scores gives an estimate of the number of correct predictions (true positives). The estimated number of errors (incorrect predictions, or false positives) is therefore the difference between the total number of images and this number. The error-rate estimate can then be defined as the ratio of the estimated number of incorrect predictions over the total number of images. This calculation can also be restricted to predictions for which confidence scores are above a given threshold, such that it allows to (i) estimate a threshold-specific error-rate (ii) find the threshold associated to a given error rate.

## 3 Results

### 3.1 Calibration at the image level

Generally, we observed that calibration (as measured by ECE) was negatively correlated with accuracy across models, for the three test datasets (Figure 3). ConvNext was the model providing the best overall performance. In particular, this model was slightly better in accuracy but much more efficient in terms of calibration (ECE of 2.37%, more than twice less than the second-best model, Swin Transformer V2, which has an ECE of 5.04%) on the Portugal dataset. In the meantime, the lightweight model, MobileNet, had bad to very bad (ECE of 34.27% on the Portugal dataset) accuracy and calibration performances.

As expected, temperature scaling allowed improving ECE values, for all models and datasets. We almost always observed a V-shape relationship between ECE and temperature, with ECE increasing quickly and by several percents around the optimum temperature value (Figure 4). This optimum temperature was generally greater than 1, suggesting that all models were initially overconfident to a greater or lesser extent. Interestingly, the V-shape curves of the different datasets overlapped well for the most accurate models (ConvNext and transformed-based models, ViT and Swin), and optimum temperature were similar across datasets. This suggested that a single optimum temperature would be sufficient to achieve efficient post-processing calibration. Indeed, using temperature scaling with temperature *T*^ˉ^, the models exhibited on average a relative reduction in ECE of 38% compared to without temperature scaling (*T* = 1) (dashed line in Figure 4).

**Figure 4:**
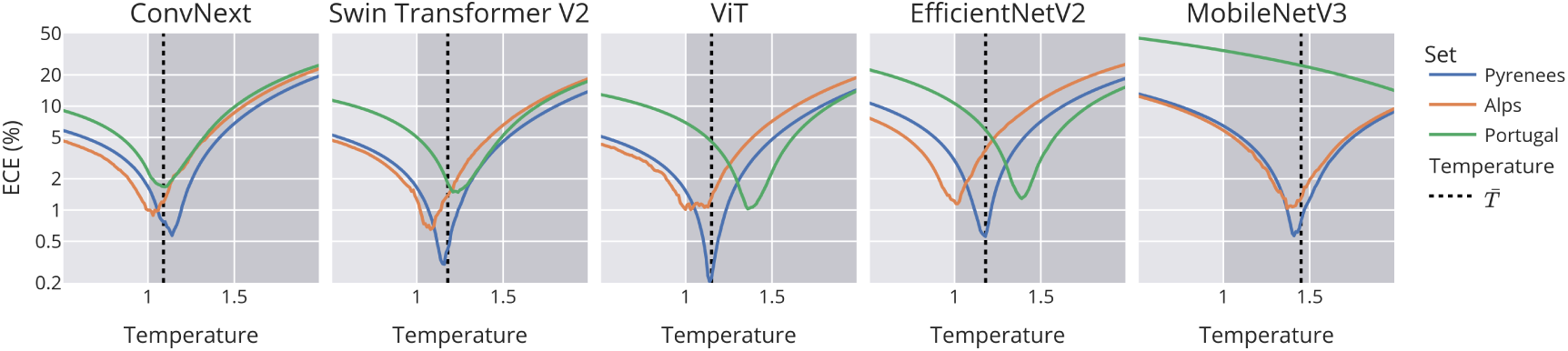
Calibration transferability using temperature scaling, at the image level. Curves of ECE values along the gradient of temperature values, for five models (panels) and three test data sets (colored curves). An optimum temperature below 1 indicates an underconfident model (light gray area), and above 1 indicates an overconfident model (dark gray area). The vertical dashed line shows *T*^ˉ^, the temperature minimizing the average ECE across the three test datasets.

### 3.2 Calibration at the sequence level

Classification accuracy was much greater at the sequence level than at the image level (Figure 5 top). This was true for all models and all datasets, with up to +10% of accuracy for MobileNetV3 on the Portugal dataset. The Average Score and Average Logit were the two best methods for maximizing accuracy, with a slight advantage for the former. Importantly, of the two aggregation methods that improved accuracy most (Average Score and Average Logit) only Average Logit provided well calibrated scores (Figure 5 bottom). The Average score was actually the worst aggregation method for calibration. Therefore, considering both accuracy and calibration metrics, the Average Logit was the best aggregation method.

**Figure 5:**
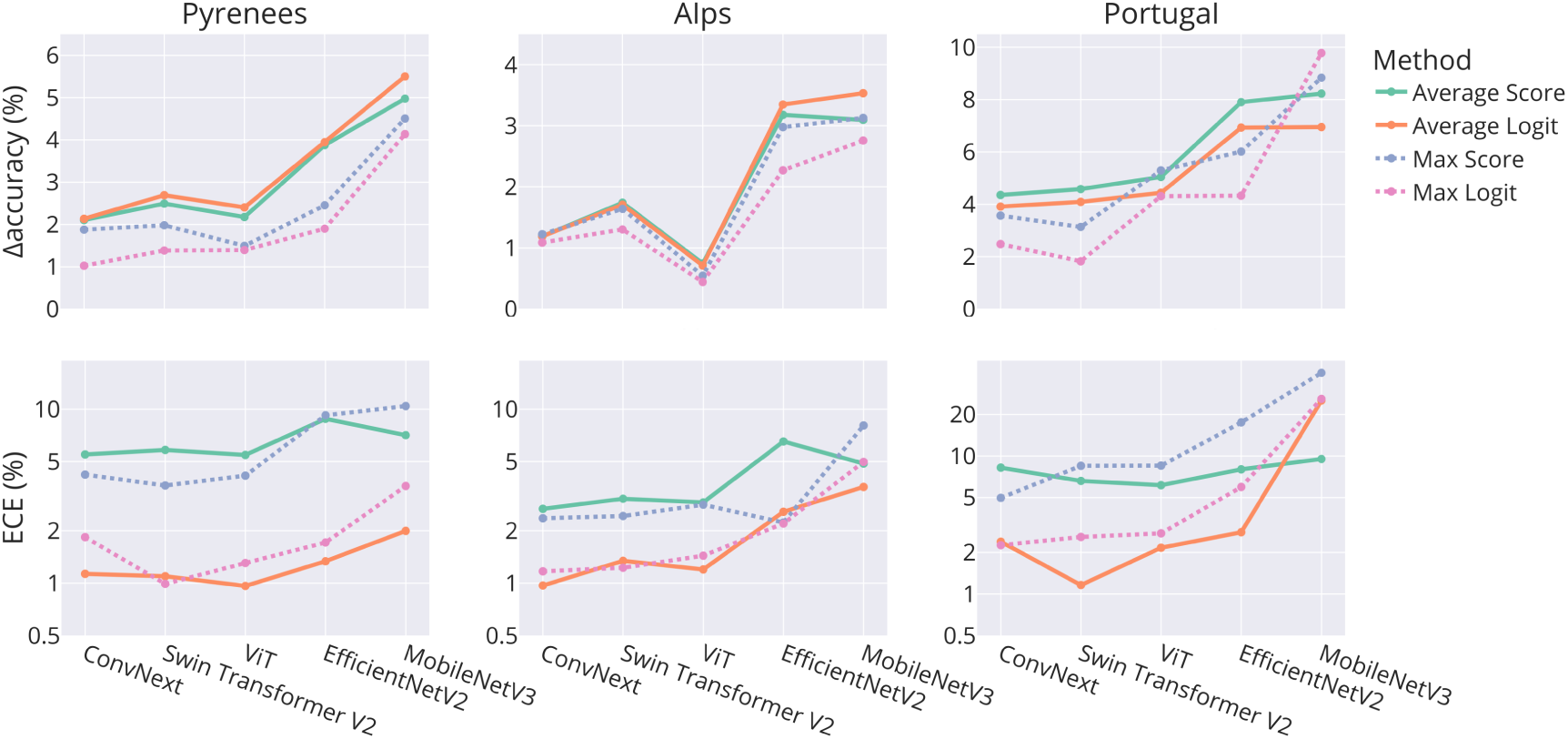
ΔAccuracy (top, the greater the better) and ECE (bottom, the lower the better) for the four aggregation methods (colored curves) and five models (x-axis) on three test data sets (three panels). ΔAccuracy is the difference between the accuracy at the sequence level and the accuracy at the image level. For the sake of clarity, solid lines represents the two best methods in terms of accuracy, Average Score and Average Logit. On the ECE plots (bottom), Average Logit outperforms Average Score in terms of calibration.

We finally studied the interplay between temperature scaling and aggregation methods. We observed that the aforementioned V-shape was more flat for the Average Score method than for the other methods (first column in Supplementary Figure 4 versus the others). This confirmed that this method was the worst method, even with temperature scaling. We also noted that the Average Logit method provided the lowest ECE values overall (1.17% on average), and thus remained the best method, with temperature scaling further improving calibration at sequence level. Finally, and as observed at the image level, a single temperature (possibly close to 1) would be sufficient to achieve good post-processing calibration with the Average Logit method.

### 3.3 Calibration use-case: controlling the number of errors along the score distribution

A practical implication of calibration is the ability to precisely estimate the number of incorrect predictions on a novel test set without labels. Here we show the impact of calibration on the quality of this estimation, using the ConvNext model (Figure 6), on the 23 353 sequences gathered after pooling the three data sets. Average Score and Max Score methods provide underconfident and overconfident scores respectively, produce inaccurate error estimations (ECE values of 6.67% and 4.38%) and under-estimate (respectively over-estimate) the number of errors by a wide margin for every threshold. For instance, with the Average Score approach, if we consider a threshold of 0.8 (often used in ecological studies, Whytock, SŚwieżewski, et al. (2021)), one would predict that the number of errors in the predictions would be approximately 500 images greater than the actual number (from 300 to 800, Figure 6). Conversely, with the Max score approach, one would predict that the number of errors in the predictions would be approximately 800 images less than the actual number (from 1200 to 400, Figure 6). Using Average Logit, the best-calibrated method without temperature scaling (ECE of 1.65%), the estimation of the number of errors is very close to the actual number. Finally, by adding temperature scaling with a temperature *T*^ˉ^ = 0.93 (ECE of 0.79%), the two curves overlap. These conclusions hold with the four other models (Supplementary Figure 5).

**Figure 6:**
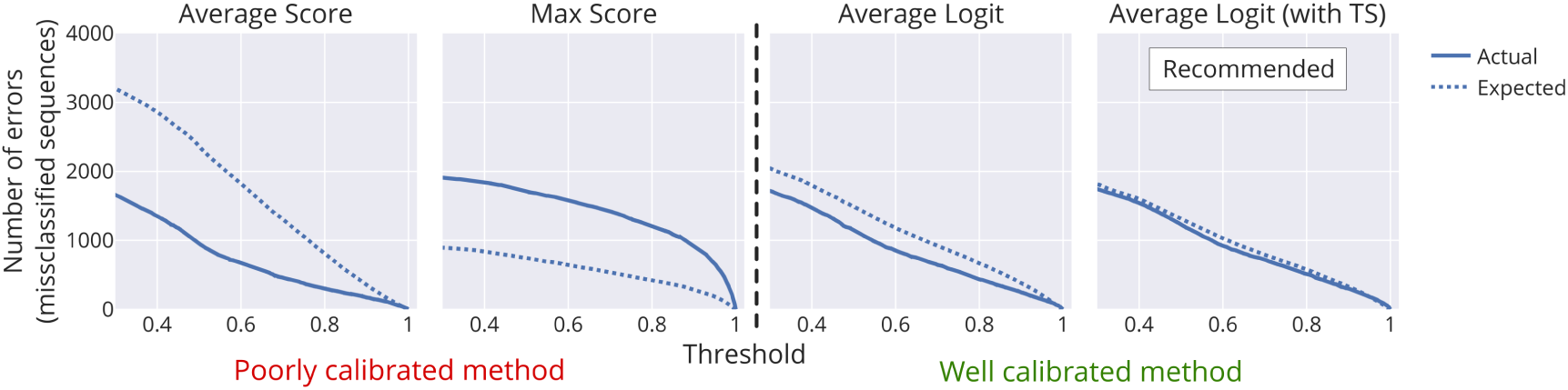
Estimation of the number of errors (i.e. misclassified sequences) when using different thresholds on the confidence scores (dash line), compared to the actual number of errors (solid line), for different aggregation methods (panels). Results shown are for the ConvNext model, the three test sets Pyrennes, Alps and Portugal being pooled together. The expected number of errors is obtained from the scores (see Material and Methods). The actual number of errors is simply the number of incorrect predictions with a score above a certain threshold, and can be known only if species labels are available. An accurate estimation of the number of errors is observed when the two lines overlap. TS means temperature scaling.

## 4 Discussion

This study assessed the calibration of confidence scores, at image and sequence level, for different deep learning models in the context of species classification in camera trap data. Using five state-of-the-art models and three out-of-sample test datasets, we showed that score calibration can vary greatly across model architectures, in a way that is consistent across test sets. Further, we showed that the different aggregation methods to obtain scores at the sequence level gave very different calibration values, and that the Average Logit method must be preferred over the others for optimizing both accuracy and calibration. Finally, we showed that temperature scaling can be used both at image level and sequence level, with a single temperature *T*^ˉ^ that do not have to vary across datasets, to further improve the calibration.

Our results about the importance of calibration in classification models are crucial for researchers and practitioners dealing with camera trap images. Firstly, we showed that confidence scores can be interpreted as probabilities only if a calibrated model is used. Secondly, we argue that the integration of machine learning predictions directly into subsequent ecological tasks can be facilitated by achieving calibration. Many ecological downstream tasks (e.g estimating occupancy, abundance or activity patterns) based on deep learning predictions use an arbitrary threshold selection (Lonsinger et al. 2023; Krivek et al. 2023) to consider that a prediction is correct, or test a series of thresholds to determine the optimal one given known species labels (Whytock, SŚwieżewski, et al. 2021; Mitterwallner et al. 2023). However, the ultimate goal of using AI is to avoid having to label images. Fortunately, we showed that calibrated models make it possible to estimate accurate error rates for any threshold, without any need of species labels. This is critical as it allows one to conduct an analysis with a known error-rate, whose level could depend on the context of the study. Calibrated models will also be central in the development of methods that can handle probabilistic uncertainties (i.e. that directly uses predictions and associated scores rather than relying on thresholding of scores) (Rhinehart et al. 2022).

Differences in models’ performance can be partly explained by model size. Indeed, we found that models with the highest number of parameters (ConvNext, ViT, SwinTransformer) gave the best accuracy and ECE values. On the other hand, the only lightweight model, MobileNet, was consistently the worst model. Despite some literature showing that neural networks can be poorly calibrated, our result shows that this is not always the case (see also Minderer et al. (2021)), and that certain families of model architectures, such as ConvNext here, are intrinsically better calibrated than others, independently of the size of the model. The calibration of each model can be further improved on each dataset using temperature scaling as post-processing function. However, determining the optimal temperature requires annotating at least a fraction of the target set of images, a step that practitioners would like to avoid if possible. Fortunately, we showed empirically with three very different datasets that the optimal temperatures are very close from one dataset to another, which suggests the generalizability of a single temperature that can be determined and fixed for future test sets. That said, we do not claim that the optimal temperatures defined in this paper can be used directly when using one of the studied architectures. Indeed, these temperatures are valid for a given training procedure (datasets, hyperparameters). In practice, it is mandatory to estimate the temperature using available test dataset(s) and subsequently maintain this temperature for deployment (since we showed it will be generalizable). This way, when the model will be used to classify new unseen data, the previously estimated temperature will ensure a better calibration of the predicted scores.

Proper model calibration at the image level is not always sufficient, as many software programs and scientific studies operate at the scale of the sequences that define the relevant ‘observations’ or ‘events’ from an ecological viewpoint. It is therefore extremely important to be able to calibrate the predictions at sequence level. For the first time, we showed that the most intuitive approach, in which scores are averaged, did not provide the best accuracy and had the worst calibration, with largely under-confident predictions. Interestingly, our findings can be confirmed by the analogy with ensemble models. These approaches use *N* models to make a prediction on *one* image, whereas we use *N* images to make a prediction with *one* model at the sequence level. Wu and Gales (2021) showed that for ensemble models, individual model calibration is not sufficient to yield a calibrated ensemble prediction, and that their own method, which is equivalent to Average Score approach also leads to under-confidence. Moreover, Rahaman and Thiery (2021) show that, thanks to this natural shift in the optimal temperature when models are ensembled, if the individual models were slightly overconfident (*T >* 1, as is often the case in deep learning) then the ensemble model was naturally calibrated (*T ∼* 1). Our results greatly support the use of the Average Logit method for aggregating individual scores at the sequence level. It shifts slightly the optimal temperature towards underconfidence, which counterbalanced the overconfident nature of deep learning networks, and resulted in sequence level prediction that are almost calibrated without post-processing. With Average Logit, it is still interesting to use temperature scaling to improve calibration as much as possible, especially given that the ECE minima are again very close to each other and allow a single temperature to be set.

In this work, we focused on temperature scaling and did not consider other methods that have been shown to sometimes improve calibration, such as label smoothing and mixup (Szegedy et al. 2015; Zhang et al. 2018). We did so because these two methods are actually debated, as several studies have showed that they can actually worsen calibration when combined with temperature scaling (Wang et al. 2023; Minderer et al. 2021). As Minderer et al. (2021) state, “label smoothing creates artificially underconfident models and may therefore improve calibration for a specific amount of distribution shift”. Label smoothing also assumes that all incorrect classes are equally likely (Maher and Kull 2021), which is obviously problematic in ecology (e.g., a wrongly predicted roe deer is much more likely to be a red deer than a wolf). Mixup also deteriorates calibration properties of networks by creating non-realistic images in the training set and leading to substantial distributional shift (Rahaman and Thiery 2021; Gawlikowski et al. 2023).

Our work concludes with some recommendations. We encourage everyone to select the architecture of their model using not only accuracy but also by calculating the ECE. Secondly, we recommend using the Average Logit method to aggregate information at sequence level, as it performs very well in terms of accuracy and calibration. Finally, to use temperature scaling and make calibration even better, the optimum temperature can be calculated on a test dataset and kept for future datasets. We acknowledge that these considerations may look too difficult to take into account for many practitioners. We therefore urge developers of camera-trap analysis software or platforms to integrate the knowledge brought by this work into their software solutions, so that it becomes immediately available the ecologists’ community and contribute to a more robust science.

## Acknowledgements

This work was granted access to the HPC resources of IDRIS under the allocation 2022-AD010113729 made by GENCI. We acknowledge the organizations and people contributing to the DeepFaune initiative.

## Author contributions

G.D., S.C.J., S.D. and V.M. conceived the ideas and designed the methodology. G.D., S.C.J. and V.M. gathered the training data. S.C.J collected the data of the Alps test set. G.D. and V.M. coded and performed the analysis. G.D. wrote the first version of the manuscript, S.C.J., S.D. and V.M. contributed critically to the drafts and gave final approval for publication.

## Data availability statement

The five trained models, all derived data used in the analysis, and the code for the inference and metric calculation are available at https://doi.org/10.5281/zenodo.10014376. The Portugal and Alps datasets are available at https://doi.org/10.15468/rah33j and https://doi.org/10.5281/zenodo.10014376. The Pyrenees dataset is available upon request only, because of the presence of a sensitive species (brown bear).

## Supplementary materials

**Figure 7:**
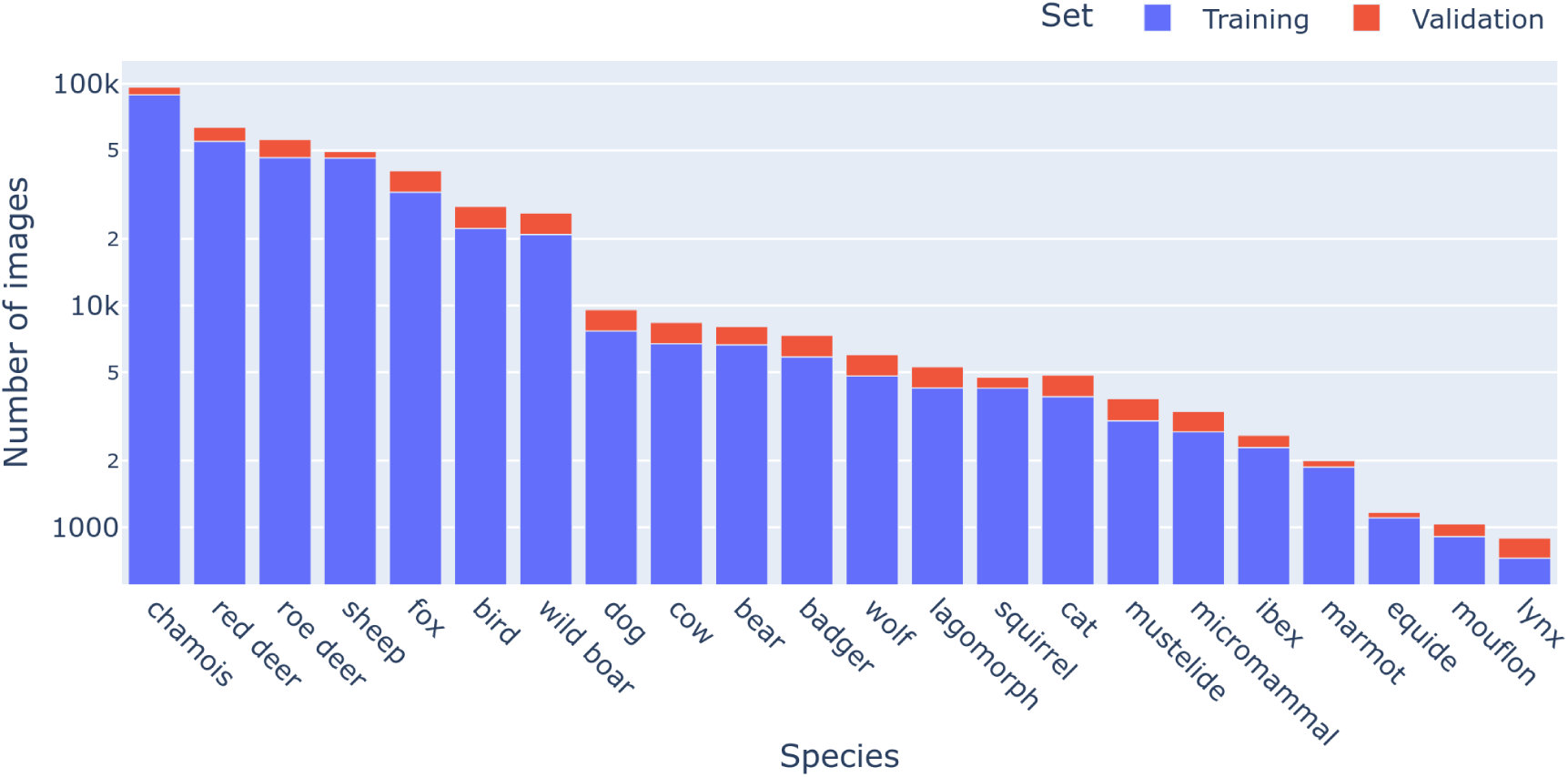
Number of images in the training and validation sets, for each species. Log scale is used to improve the readability of the rarer classes.

**Figure 8:**
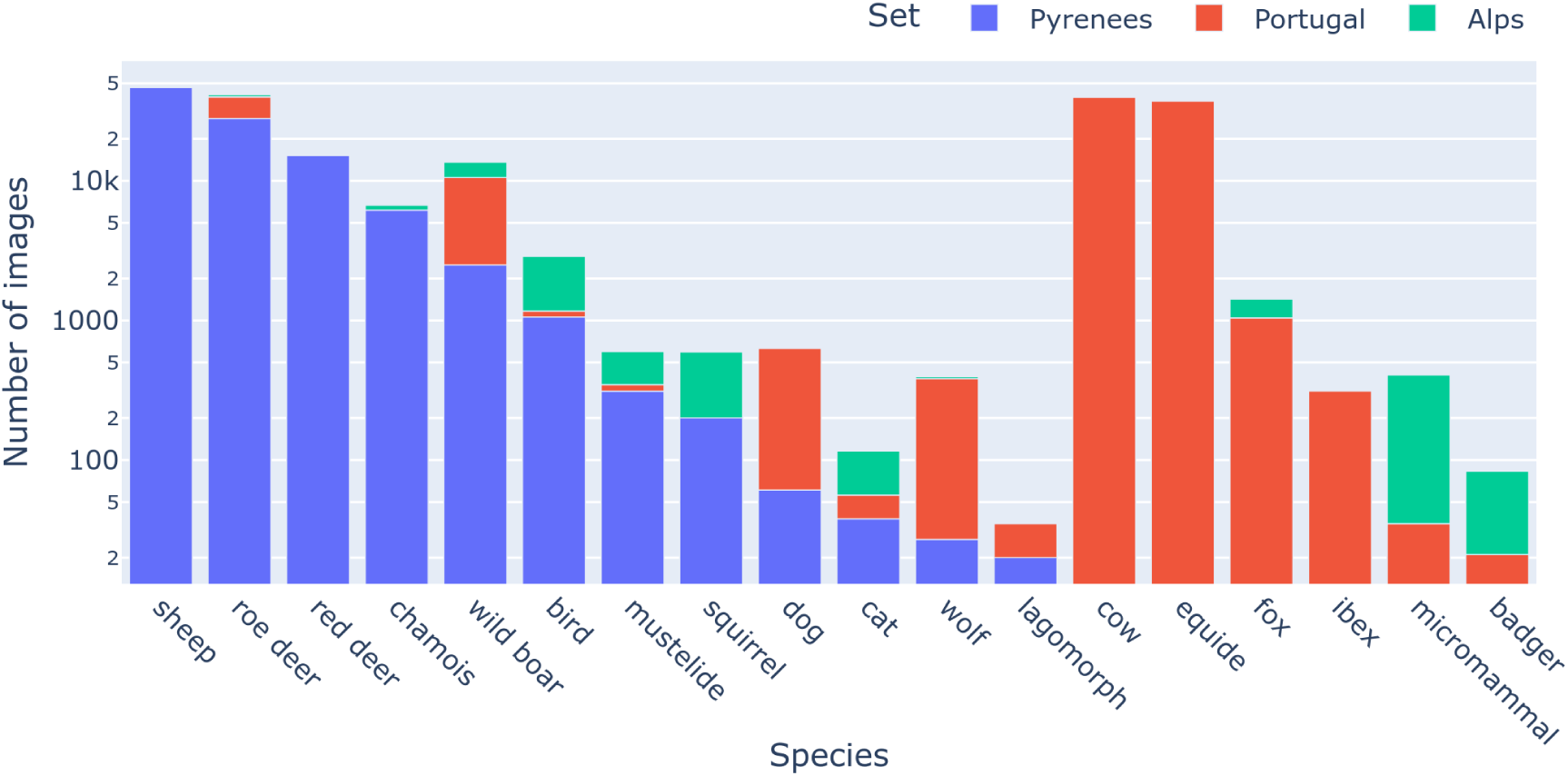
Number of images in the three out-of-sample datasets, for each species. Log scale is used to improve the readability of the rarer classes.

### 4.1 Results

**Figure 9:**
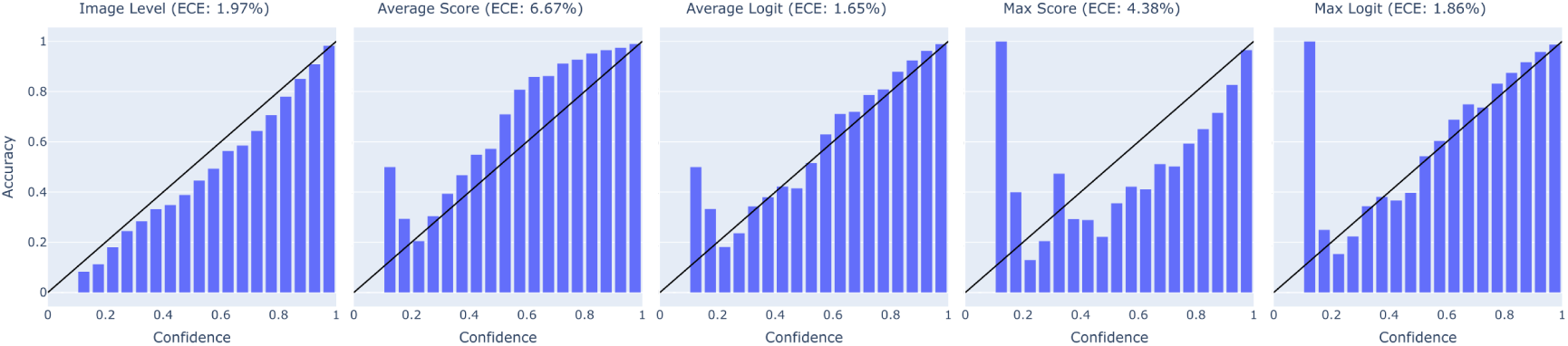
Reliability histogram of the ConvNext model, using the 3 test sets pooled together, and without temperature scaling.

**Figure 10:**
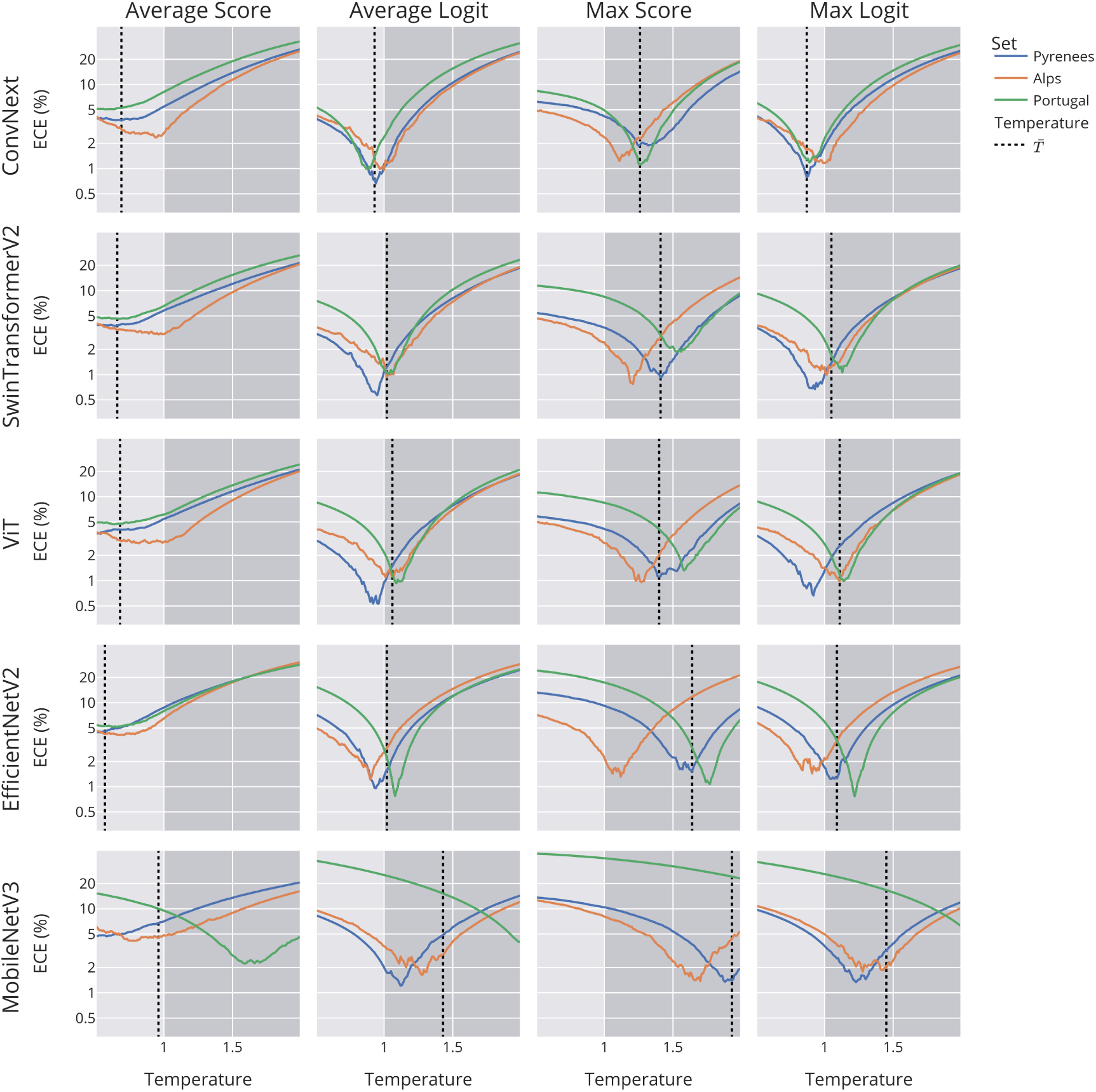
Calibration transferability using temperature scaling, at the sequence level. Curves of ECE values along the gradient of temperature values, for the four aggregation methods (columns), the five models (rows) and the three test datasets (colored curves). Light/gray area and dashed line defined as in the main text, Figure 4.

Looking at *T*^ˉ^ and the optimum temperature of each set (minima and dashed lines on Figure 10), it can be noted that using the Average Score methods led models towards underconfidence, whereas using the Max Score methods led them towards overconfidence. This observation is also visible in the reliability histogram, as shown in Supporting Information (Supplementary Figure 3).

**Figure 11:**
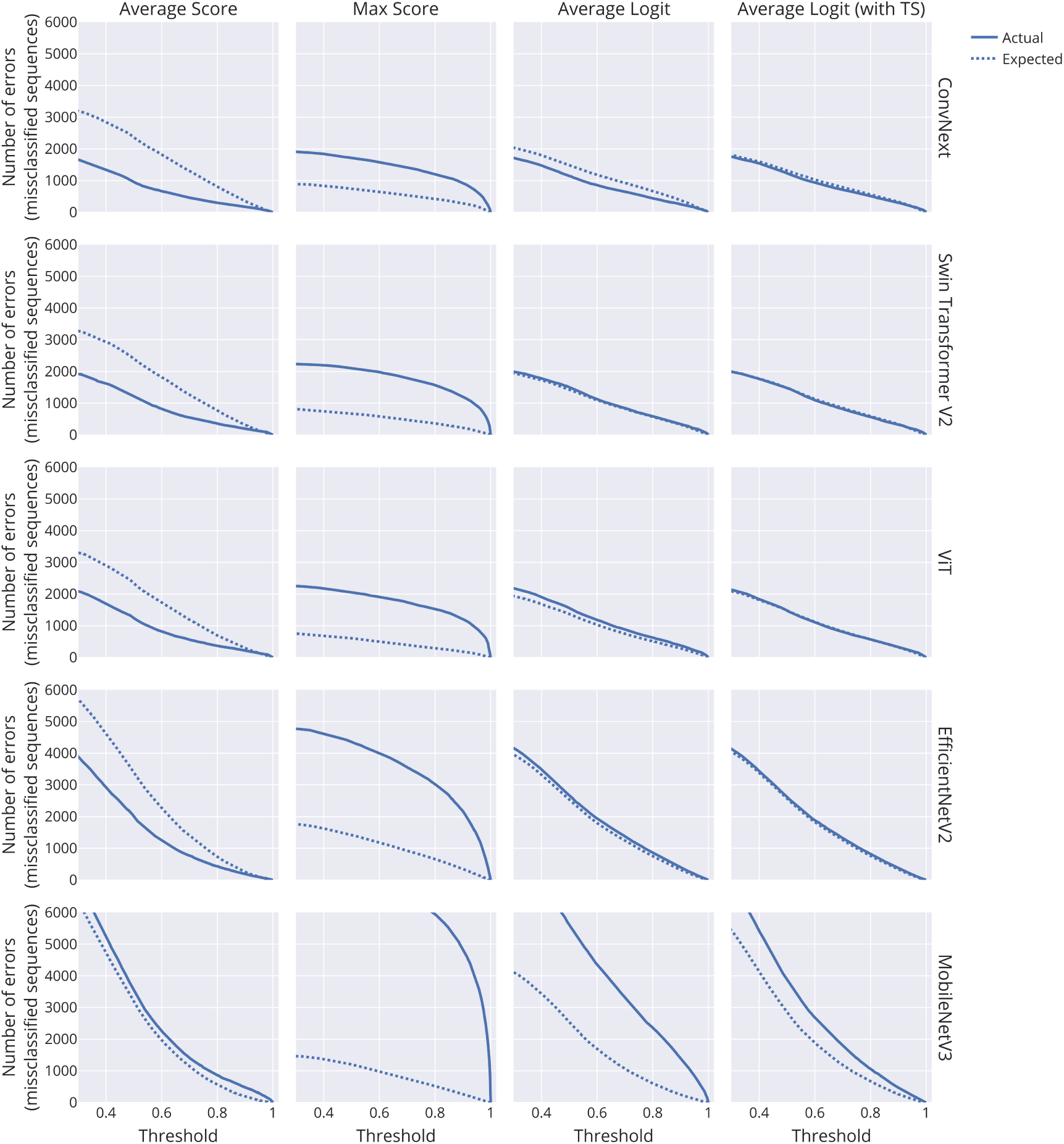
Estimation of the number of errors (i.e. misclassified sequences) as a function of the confidence score (dashed line), compared to the actual number of errors (solid line), for different methods. The three test sets are pooled together.

